# Early Ictal Recruitment of Midline Thalamus in Mesial Temporal Lobe Epilepsy

**DOI:** 10.1101/647586

**Authors:** Andrew Romeo, Alexandra T. Issa Roach, Emilia Toth, G. Chaitanya, Adeel Ilyas, Kristen O. Riley, Sandipan Pati

## Abstract

The causal role of midline thalamus in the initiation and early organization of mesial temporal lobe seizures is studied. Three patients undergoing stereoelectroencephalography were enrolled for the placement of an additional depth electrode targeting the midline thalamus. The midline thalamus was recruited in all three patients at varying points of seizure initiation (0-13 seconds) and early propagation (9-60 seconds). Stimulation of either thalamus or hippocampus induced similar habitual seizures. Seizure-induced in the hippocampus rapidly recruited the thalamus. Evoked potentials demonstrated stronger connectivity from the hippocampus to the thalamus than in the opposite direction. The midline thalamus can be within the seizure initiation and symptomatogenic circuits.

## Introduction

Resective epilepsy surgery can be curative and potentially life-saving if seizure freedom is sustained over an extended period^1^. Despite advances in imaging and surgical techniques, the surgical outcome of drug-resistant temporal lobe epilepsy (TLE) has plateaued, with only 40-60% patients attain long term seizure freedom following anterior temporal lobectomy (ATL)^2^. One prevailing hypothesis of seizure recurrence post-ATL is that a subset of patients have an underlying epileptogenic network extending beyond the temporal “focus” and may include subcortical structures including the thalamus^3, 4^. Multiple structural and functional MRI studies have demonstrated the persistence of thalamotemporal connectivity as a predictor of seizure recurrence and a marker of poor prognosis following ATL^3, 5^. In TLE, thalamic subnuclei play a diverse role in the initiation and propagation of seizures. The anterior nucleus has gained substantial interest recently from the success of deep brain stimulation (DBS) therapy in focal epilepsy, but the pulvinar, mediodorsal, and centromedian nuclei have all been studied in human epilepsy^4, 6, 7^. In preclinical models of TLE, the midline thalamus (reuniens and rhomboid nuclei) have – a) the majority direct thalamic input to the hippocampus with reciprocal connectivity to other limbic cortices (amygdala, anterior cingulate, medial prefrontal cortex); b) is involved in the initiation of limbic seizures; c) has excitatory influence on the epileptogenic hippocampus; and d) its inhibition can reduce induced seizures from the hippocampus^4, 8-11^. A human electrophysiological study replicating some of these preclinical findings is lacking. In this first-in-man study that incorporates direct electrophysiological recordings from the midline thalamus, we hypothesize that mesial temporal lobe seizures recruit the midline thalamus early at seizure onset and before the clinical manifestation.

## Methods

### a) Patient selection and data acquisition

Three consecutive patients with suspected TLE undergoing stereo EEG investigation at a level-IV center were enrolled prospectively for placement of an additional depth electrode targeting the midline thalamus. The local Institutional Review Board approved the study protocol, and written consent was obtained before research implantation in the thalamus. Percutaneous stereotactic placement of Depthalon (PMT) electrodes was performed using the ROSA robot (Medtech). A tailored implantation strategy for each patient was designed based on the preoperative hypothesis of the underlying epileptic network. Video EEG was sampled at 2048 Hz using Natus Quantum (Natus Medical Incorporated, Pleasanton, CA). Seizures were defined as the emergence of rhythmic or repetitive spikes lasting more than 10 s with or without clinical manifestations, and seizure onset was defined as the earliest occurrence of rhythmic or repetitive spikes. Board-certified epileptologists annotated the seizures, and the seizure onset zone was localized by integrating anatomo-electroclinical features in a multidisciplinary conference.

### b) Targeting the midline thalamus

For the purpose of this paper ventral midline thalamus (VMT) and midline thalamus are interchangeable. There are <u>VMT nuclei in both the left and right thalami on either side of the third ventricle.</u> The VMT nuclei near the massa intermedia was targeted for implantation. The widest midline nuclear segment is anterior near the massa intermedia where the reuniens and central median nuclei are located. The inferior and slightly posterior relationship with the anterior thalamic nucleus provided further anatomic confirmation. The depth was planned to abut the third ventricle. A trajectory targeting the frontal operculum/insula was extended to end at the thalamic target. The implantation strategy allowed research recording from the thalamus without placing an additional depth electrode. A lateral trajectory with an entry point in the frontal operculum was chosen to avoid traversing the ventricle. The electrode passed through the mediodorsal nucleus before terminating in the ventral midline region (Fig1A, supplementary Figure 3). Post-implantation CT imaging was co-registered with preimplantation T1-weighted 3 T MRI to affirm accurate electrode localization^12^.

**Figure 1:**
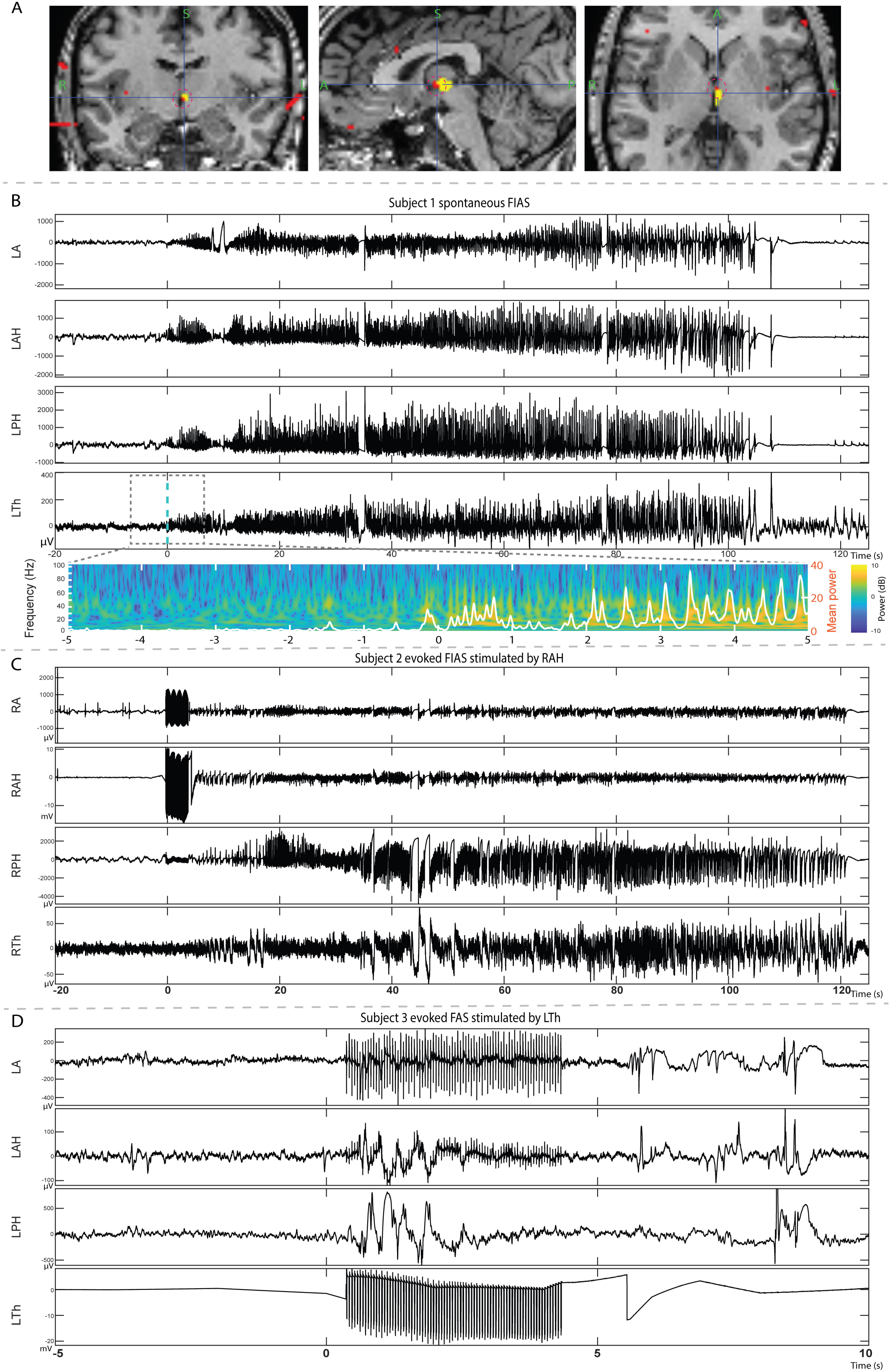
A-Post-implant CT brain coregistered with MRI to demonstrate depth electrodes (highlighted with red dots) targeted towards the midline thalamus (highlighted with yellow dots) B-Stereo EEG recording demonstrating spontaneous seizure (FIAS) localized to left amygdala (LA), anterior and posterior hippocampus (LAH, LPH), and midline thalamus (LTh). EEG changes low amplitude fast activity (LAFA). Included below is the time-frequency decomposition (1-100Hz) of ictal thalamogram. Note the increase in power above 15 Hz following ictal recruitment. C-Stereo EEG recording demonstrating induced seizure (FIAS) localized to the right amygdala (RA), anterior and posterior hippocampus (RAH, RPH), and midline thalamus (RTh). Electrical stimulation was applied to RAH. D-Stereo EEG recording demonstrating evoked seizure (FAS) localized to left amygdala (LA), anterior and posterior hippocampus (LAH, LPH). Electrical stimulation was applied to left midline thalamus (LTh). Abbreviation-FIAS: Focal impaired awareness seizure; FAS-Focal aware seizure

### c) Direct Cortical Stimulation of the SEEG electrodes

Direct electrical stimulation (Nicolet Cortical Stimulator) of the SEEG electrodes was performed to induce seizures on the last two days of SEEG investigation after antiepileptic drugs were resumed. Bipolar, charge balanced biphasic stimulation at 50 Hz was performed with pulse width 400 us, burst duration 5 seconds and current between 4-5 mAmp.

### d) Corticocortical evoked potentials (CCEP)

CCEP mapping was performed to establish the causal relation (“effective connections”) between the midline thalamus and hippocampus^13^. Bipolar stimulation of each pair of adjacent electrodes was performed at 1 Hz with electrical current 3mA, pulse width 300 us for 40 trials. The recordings were detrended (0.5 s) during pre-processing and bipolar montage was used to analyze. The maximal amplitude of every evoked response (10 to 100 ms after stimulation) was z-score transformed using the baseline (−100 to −2 s) mean and standard deviation. Epochs without higher than 3 z-score maximal value were excluded from the average^13^.

### e) Time-frequency decomposition of thalamogram

Preprocessed data was S-transformed between 1-100 Hz, and average power was plotted with an overlaid white line^14^.

## Results

### a) Ictal recruitment of midline thalamus

Demographic, seizure and outcome information regarding the three patients are summarized in Table 1A. Subject 1 had 12 seizures (focal impaired awareness seizure-FIAS)^15^ with onset in the left amygdala-hippocampus (Amy-Hippo) and rapid (either simultaneously or within seconds after cortical onset) recruitment of the left thalamus (Fig1B). The electrographic onset pattern consisted of low amplitude fast activity (LAFA) in the Amy-Hippo and thalamus (Table 1B, Supplementary Fig.1A). Subject 2 had 6 seizures (FIAS) with heralding spike and LAFA localized to the right hippocampus and the ictal activity propagated to ipsilateral thalamus before clinical onset within several seconds(Table1B, Supplementary Fig1B). Subject 3 had over 40 seizures (focal aware seizure-FAS and electrographic) with hypersynchronous spikes localized to the left Amy-Hippo, with late recruitment of the thalamus immediately before or after clinical onset (Supplementary Fig1C).

**Table 1-A:** Demographics, and details of seizure semiology, stereo EEG investigation of three participants.

| Subject Number | Age (yrs) | Gender/handedness | Age of Onset (years) | Reported Seizure semiology | MRI | Thalamic implant side | SEEG implant strategy | Recorded seizures | SOZ localization | Post SEEG therapy and outcome |
| --- | --- | --- | --- | --- | --- | --- | --- | --- | --- | --- |
| 1 | 22 | F/L | 13 | a. arousal from sleep →oro manual automatism →head version to right →right upper extremity clonic activity<br>b. facial paresthesia →oro manual automatism →dialeptic (rare events) | Nonlesional | L | L=10<br>R=9 | FIAS=12 | Left amygdala-hippo and left midline Thalamus* | deferredRNS (left hippo depth +left subtemporal strip) after a trial of Epidiolex |
| 2 | 39 | F/L | 15 | a. epigastric rising sensation→metallic taste<br>b. epigastric rising sensation→oro manual automatism →dialeptic | Nonlesional | R | R=8 | FIAS =6 | Right hippo-amygdala | Right ATL. Histopathology CA4 neuronal loss and gliosis(ILAE Type3). Seizure free at 3 months |
| 3 | 42 | F/ambidextrous | 8 | a. olfactory hallucination (“stinky smell”)→ blurry vision → grabs nose with right hand<br>b. olfactory hallucination (“stinky smell”)→grabs nose with right hand→dialeptic | L and R hippo sclerosis, global cortical atrophy | L | L=11<br>R=7 | FAS=28<br>E= 12 | Left hippo-amygdala | left amygdala hippo LITT#<br>Seizure free at 2 months |

\* SEEG
recording from the thalamus was for research purpose only and hence RNS was not placed in the thalamus. # significant cognitive impairment precluded from alternative therapies like RNS or open ATL. Abbreviations: hippo= hippocampus; F=Female; R/L=Right/left; FIAS= Focal impaired awareness seizure; FAS= Focal aware seizure; E=Electrographic; LITT- Laser interstitial thermal therapy; RNS= Responsive Neurostimulation

**Table 1-B:** Details of electrographic changes with spontaneous and evoked seizures.

| Subject Number | Seizure onset zone (SOZ) | Recorded Seizure subtype | Ictal EEG in SOZ | Ictal EEG in Midline thalamus | Clinical onset latency after electrographic onset (in secs) – Mean, (Range in secs) | First unequivocal ictal EEG changes in the thalamus-electrographic onset of seizure (Range in secs). Data for clinical seizures only |
| --- | --- | --- | --- | --- | --- | --- |
| 1 | Left hippo, AMY, midline thalamus | FIAS | LAFA | LAFA | 5 secs (3-8 secs) | 0 to 2 secs |
| 2 | Right hippo | FIAS | Heralding spikes→ LAFA | LAFA | 21 secs (11-72 secs) | 4 to 13 secs |
| 3 | Left hippo, AMY | FAS | Hypersynch spikes→ LAFA | Rhythmic spikes | 16 secs (9-64 secs) | 9 to 72 seconds |

Abbreviations: LAFA= Low amplitude fast activity; SOZ= seizure onset zone; Hypersynch= hypersynchronous; Hippo= Hippocampus; AMY- Amygdala; FAS= Focal Awareness Seizure; FIAS= Focal Impaired Awareness Seizure

### b) Electrical Stimulation(ES)of thalamus induced habitual seizures

For all three subjects, ES of thalamus induced habitual seizures without any ictal EEG changes in the thalamus and the EEG changes in the Amy-Hippo were variable in morphology (Table1C). For the subject 1, the induced ictal EEG consisted of hypersynchronous spikes in the Amy-Hippo while for Subject 3 there was a burst of spike-wave complexes restricted in the Amy-Hippo only (Fig1D). Subject 2 had FSA and had no EEG changes. Interestingly, for all the subjects stimulation of hippocampus induced habitual electroclinical seizures with rapid propagation to the thalamus (Fig 1C, Supplementary Fig 2A-C).

**Table 1-C:** Details of evoked seizures following direct electrical stimulation of hippocampus and thalamus.

| Subject Number | Stimulation of hippocampus | Stimulation of midline thalamus |
| --- | --- | --- |
| 1 | habitual FSIA (# b in Table 1A) with rapid recruitment of ipsilateral thalamus | Habitual FSIA (#b in Table 1A) without ictal EEG changes in the thalamus. EEG in Amyg-hippo showed low amplitude hypersynchronous spikes |
| 2 | habitual FSIA (#b in Table 1A) with rapid recruitment of ipsilateral thalamus | habitual FSA (#a in table 1A) without ictal EEG changes in the thalamus, Amyg-hippo |
| 3 | habitual FSA (#a in Table 1A) with recruitment of ipsilateral thalamus | habitual FSA ( #a in Table 1A) with short lasting epileptiform activities in the amygd-hippo but no ictal changes in the thalamus |

FAS= Focal Awareness Seizure; FIAS= Focal Impaired Awareness Seizure

### c) Variable reciprocal connectivity between midline thalamus and epileptogenic hippocampus

CCEPs demonstrated bi-directional but asymmetrical connectivity between the midline thalamus and anterior hippocampus (stronger connectivity from the hippocampus to the thalamus than vice versa) (Fig.2). The CCEP_hippo→thal_ was higher for Subject 1 than in Subject 2 (amplitude z scores - 34.1 vs. 6.3) (Fig.2D,E). In Subject 3, a single pulse stimulation of anterior hippocampus induced a habitual seizure and hence CCEPs was not obtained.

**Figure 2:**
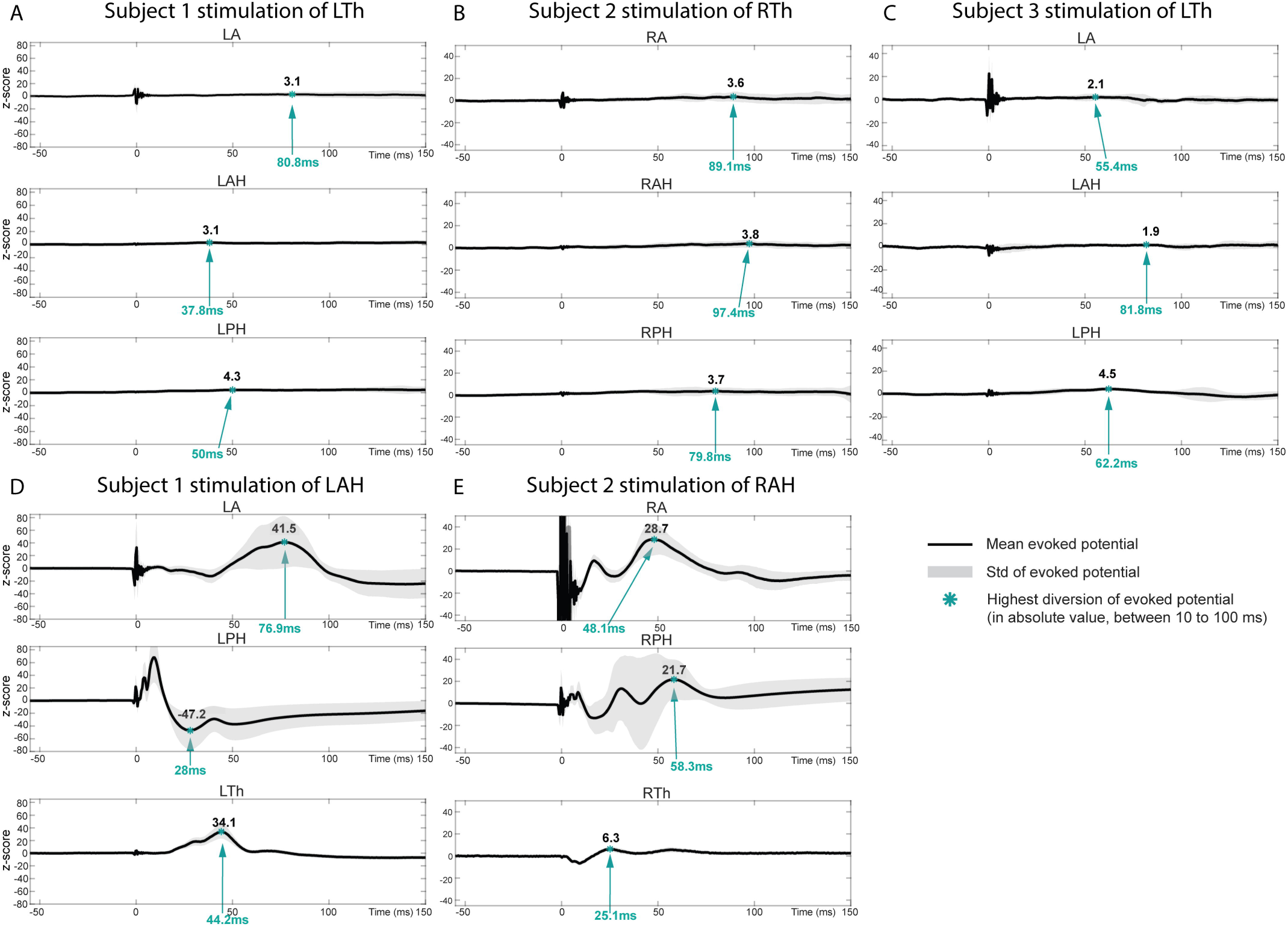
Corticocortical evoked potential (CCEP) for subject 1-3. The results of 1Hz stimulation of midline thalamus(Th) (A-Subject1; B-Subject2; C-Subject 3) and anterior hippocampus(AH) (D-Subject 1; E-Subject 2,). PH-Posterior hippocampus; A-Amygdala.

## Discussion

Increasingly studies have demonstrated the distributive nature of the seizure focus in TLE, with connections of the mesial temporal regions to distant sites like the insula or orbitofrontal regions^16, 17^. Identifying the remote ictogenic sites is critical as ATL in these “Temporal-plus” epilepsies can only achieve suboptimal seizure control^16, 17^. Here, we demonstrate in humans the midline thalamus as another remote structure that can be within the neural circuit associated with initiation and propagation of mesial temporal onset seizures. The study highlights the temporal variability in the ictal recruitment of midline thalamus. The strength of connectivity between the hippocampus and the midline thalamus varied between the patients. The stronger connectivity from the hippocampus to the thalamus in Subject 1 might explain the fast recruitment of thalamus at seizure onset.

The connectivity of the human midline thalamus is unknown, but the nucleus reuniens in rodents has strong excitatory projections to CA1 and mesial prefrontal cortex^9, 11^. The excitatory input of the midline thalamus to the hippocampus can potentially alter excitability and generate seizure from the hippocampus^9^. In all three subjects we were able to induce the habitual seizure by stimulating the thalamus, but ictal EEG changes in the cortex were variable. One explanation for lack of ictal EEG changes could be that stimulation of the thalamus resulted in excitation of the interconnected cortical regions that are involved in initiation of seizures but are under (or partially) sampled with SEEG. Although imaging studies have demonstrated the presence of thalamic hub in prognostication of ATL at a group level^3, 5^, electrophysiological studies including direct cortical stimulation can clarify the role of the thalamus in ictogenesis at an individual level. A smaller number of patients and the lack of surgical outcome are limitations in this exploratory study.

## Conclusion

Mesial temporal lobe seizures can recruit midline thalamus at seizure onset, and habitual seizures can be replicated by stimulating the thalamus. Unlike the anterior thalamus that is implicated in propagation of temporal lobe seizures, the midline thalamus can be within the seizure onset network in TLE that needs to be clarified in a larger cohort. With the widespread acceptance of robot-assisted sEEG investigation to localize epilepsies, electrophysiological sampling from the thalamus is feasible and maybe justifiable in selected patients where the demonstration of early thalamic recruitment can be used to improvise therapies (-like implanting responsive neurostimulation in the thalamus^18^) that complement resection.

## Acknowledgments

All authors greatly appreciate the patients that have participated in this study. SP gratefully acknowledge the support from the USA National Science Foundation (NSF RII-2FEC OIA1632891).

## Author Contributions

S.P. conceived the study plan and was involved in study design, data acquisition, analysis and interpretation. A.T.I.R, E.T., G.C, A.I. performed data analysis and interpretation. A.R., K.O.R. were involved in data acquisition. All authors were responsible for drafting the manuscript and gave approval for the final manuscript.

