## Supplemental Fig for "Early Ictal Recruitment of Midline Thalamus in Mesial Temporal Lobe Epilepsy"

**Supplementary Figures**

Fig 1 : Stereo EEG recordings demonstrating spontaneous seizures. EEG windowed between 1-100 Hz . * represent electrographic onset , ** clinical onset. Bipolar recordings from the midline thalamus is highlighted in red. A : Subject 1- FIAS localized to left amygdala , anterior and posterior hippocampus and midline thalamus (LT1-2). Time base 15 mm/sec

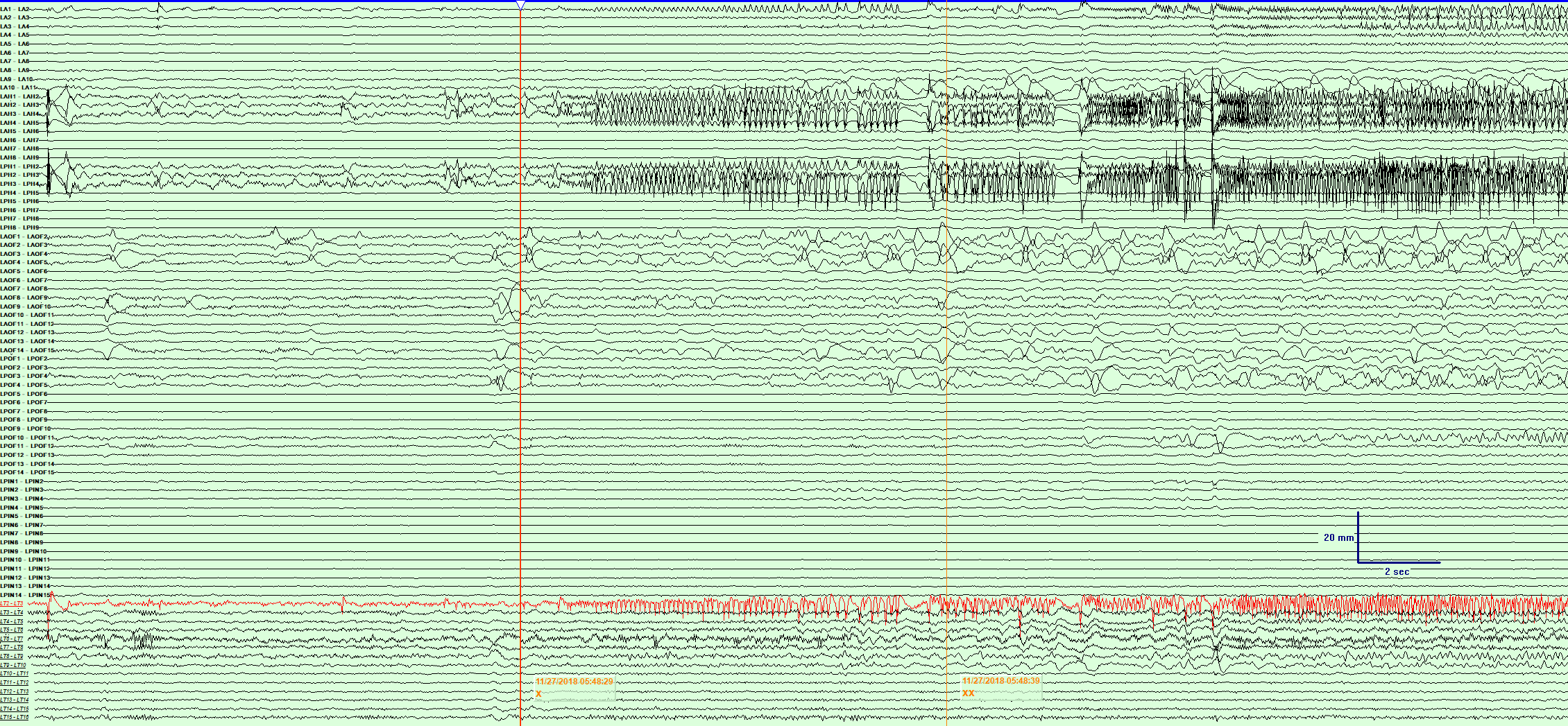

B : Subject 2- FIAS localized to right amygdala , anterior and posterior hippocampus. Note ictal spread to the midline thalamus (RT1-2).

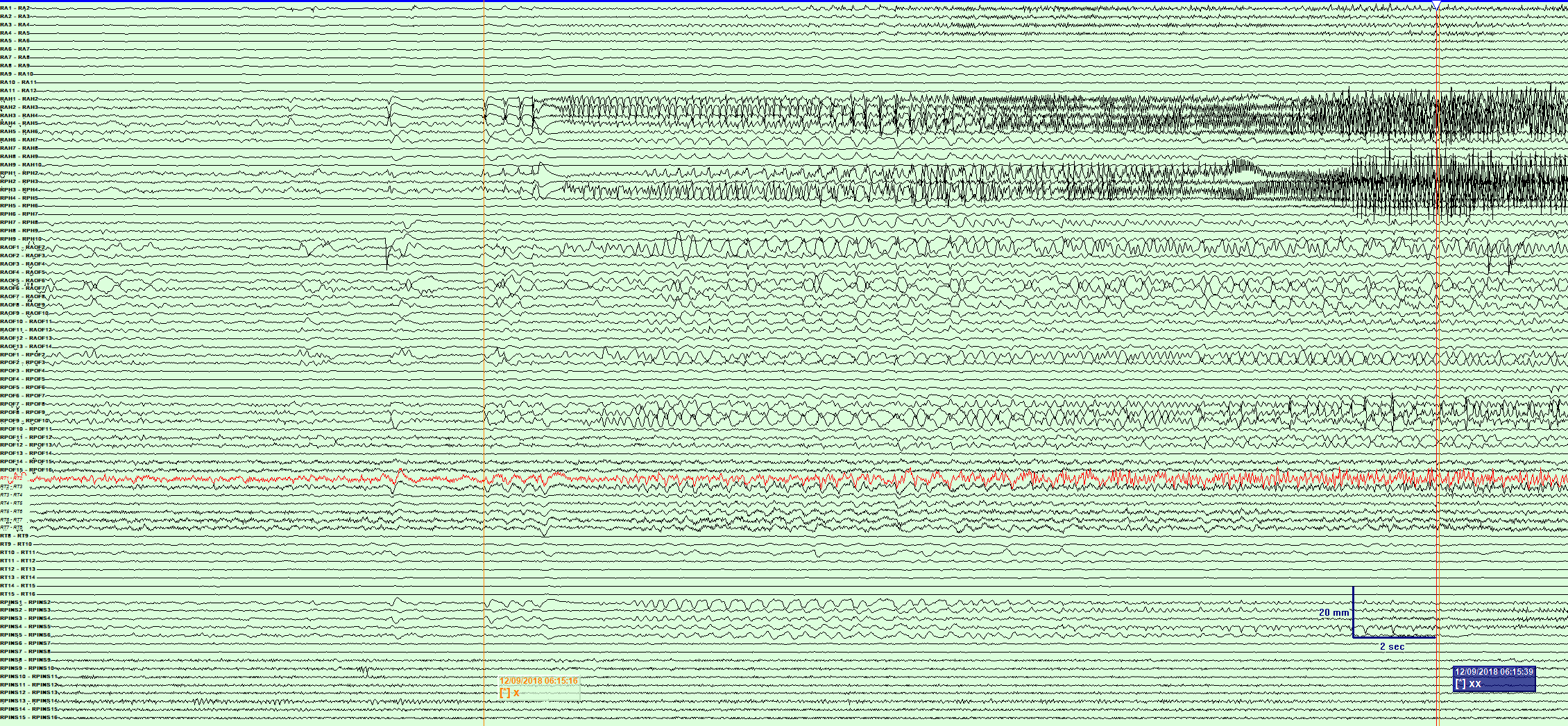

C : Subject 3- FAS localized to left amygdala , anterior and posterior hippocampus. Note ictal spread to the midline thalamus (LT1-2) late during behavioral onset (marked **).

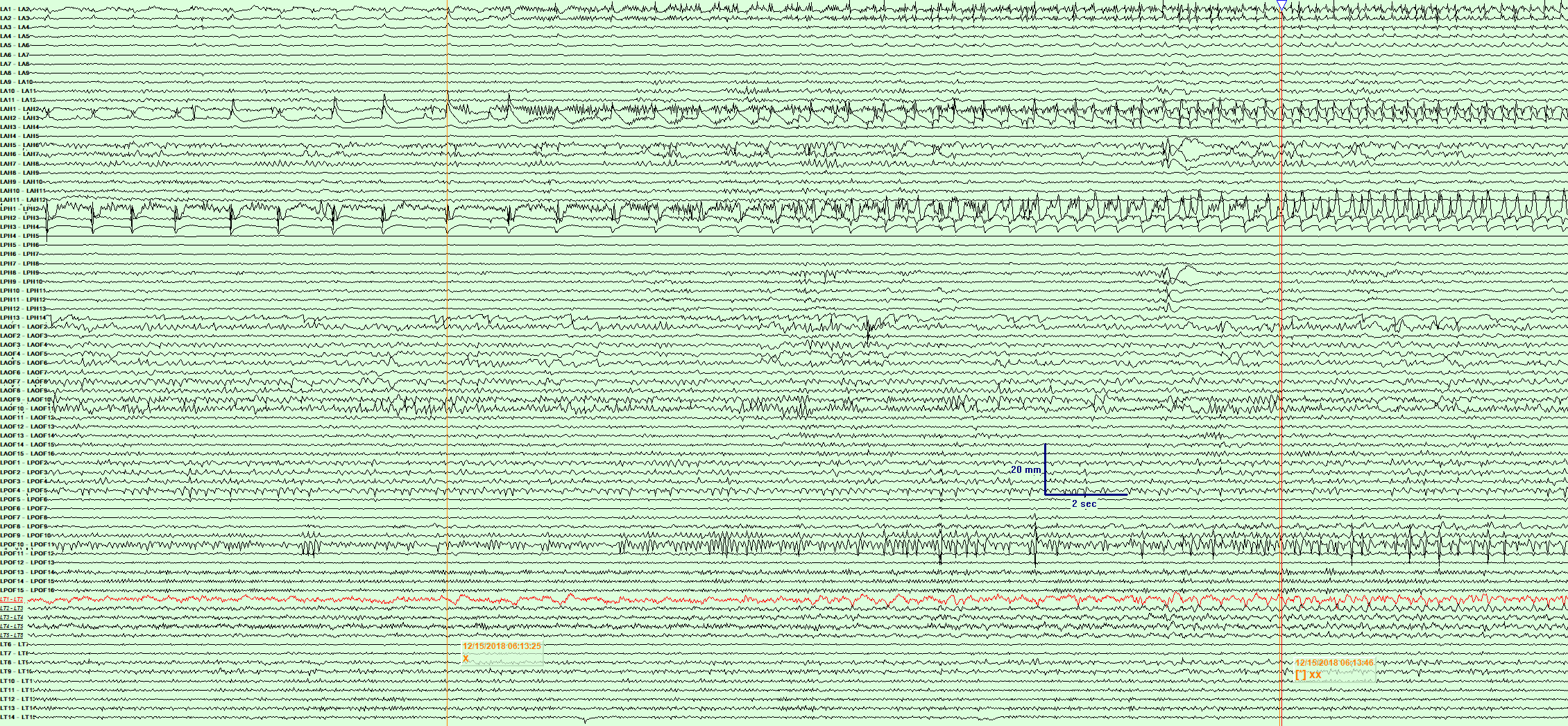

Fig 2 : Stereo EEG recordings demonstrating induced seizures. Direct electrical stimulations performed in the anterior hippocampus. EEG windowed between 1-100 Hz. Time base 15 mm/sec (except figure C) . A : Subject 1- Induced FIAS localized to left amygdala , anterior and posterior hippocampus and midline thalamus (LT1-2).

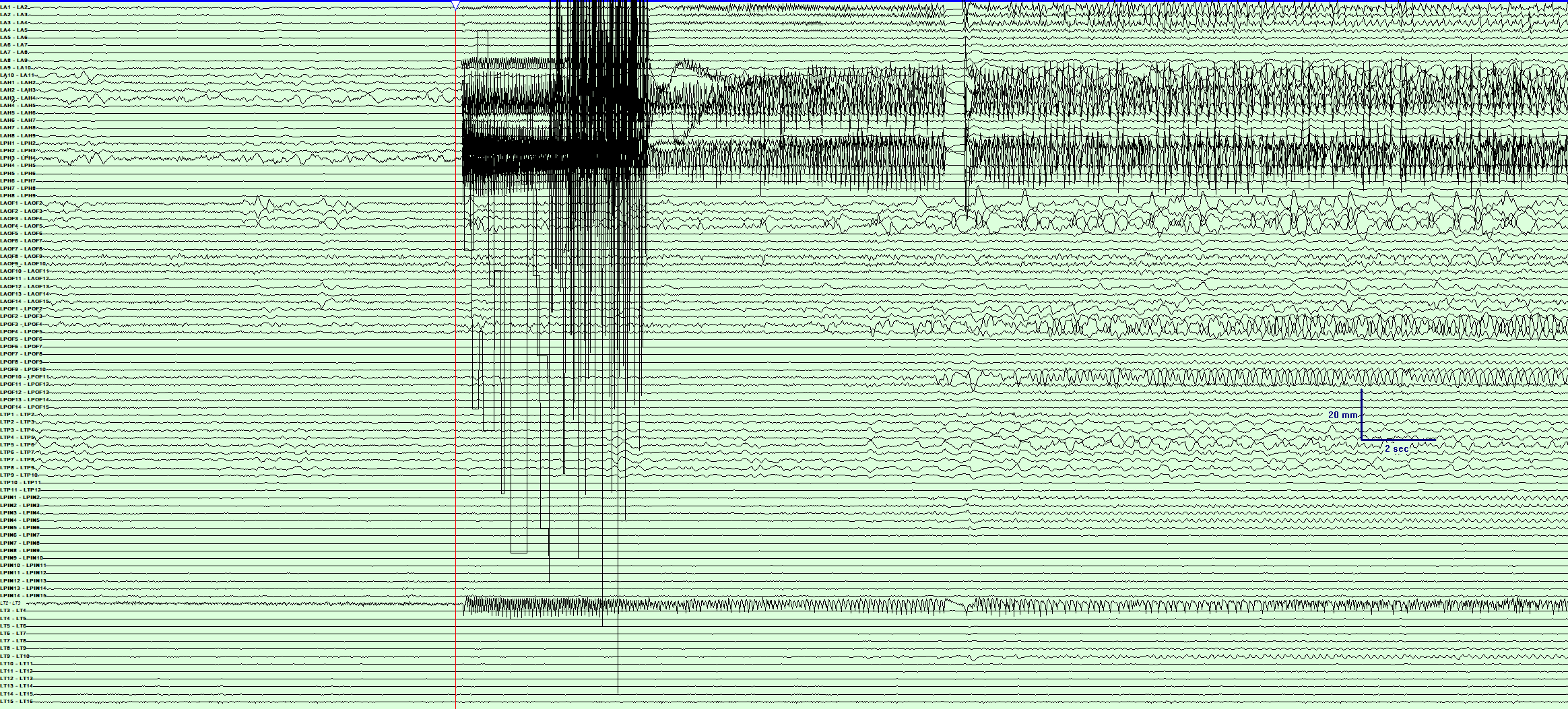

B : Subject 2- Induced FIAS localized to right amygdala , anterior and posterior hippocampus. Note ictal spread to the midline thalamus (RT1-2).

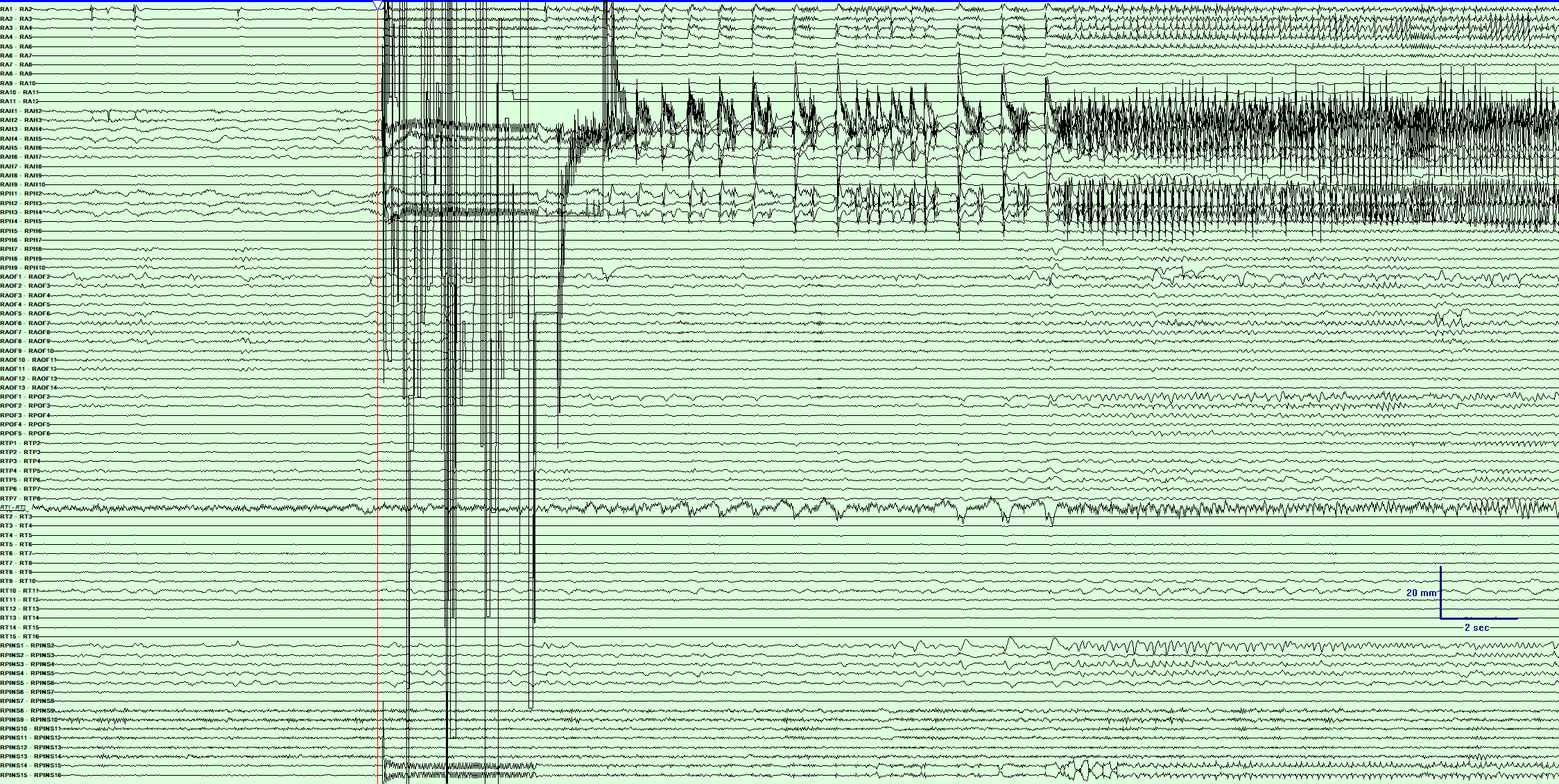

C : Subject 3- Induced FAS localized to left amygdala , anterior and posterior hippocampus. Note late ictal spread to the midline thalamus (LT1-2). Note- the time base is increased to 30 mm/sec in order to demonstrate the late recruitment of midline thalamus

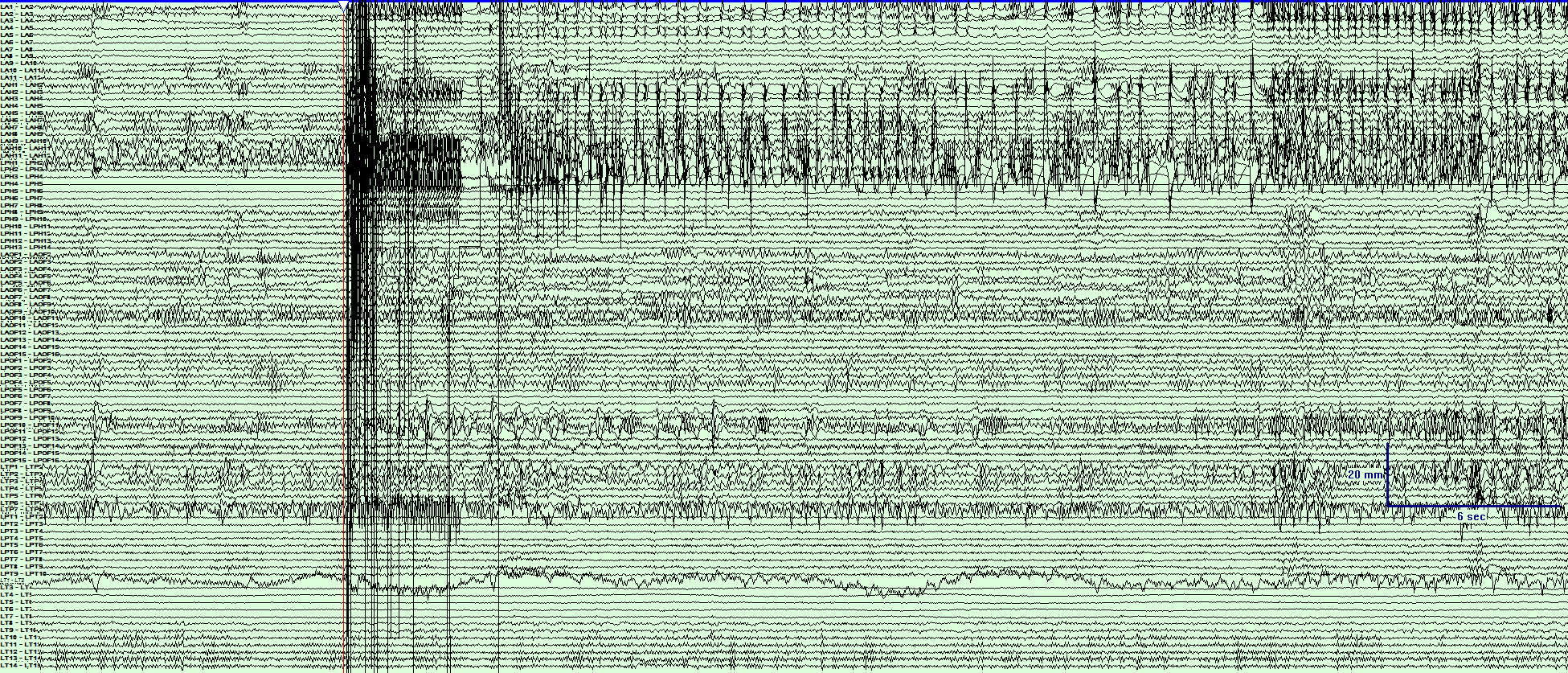

Figure 3: - Post-implant CT brain coregistered with MRI to demonstrate depth electrodes (highlighted with white dots) targeted towards the midline thalamus (highlighted with red dot). For each subject coronal (above) and axial views (below) provided. Subject 2 and 3 are provided below. Subject 1 is in Fig 1A (main paper).

**Subject 2	Subject 3**

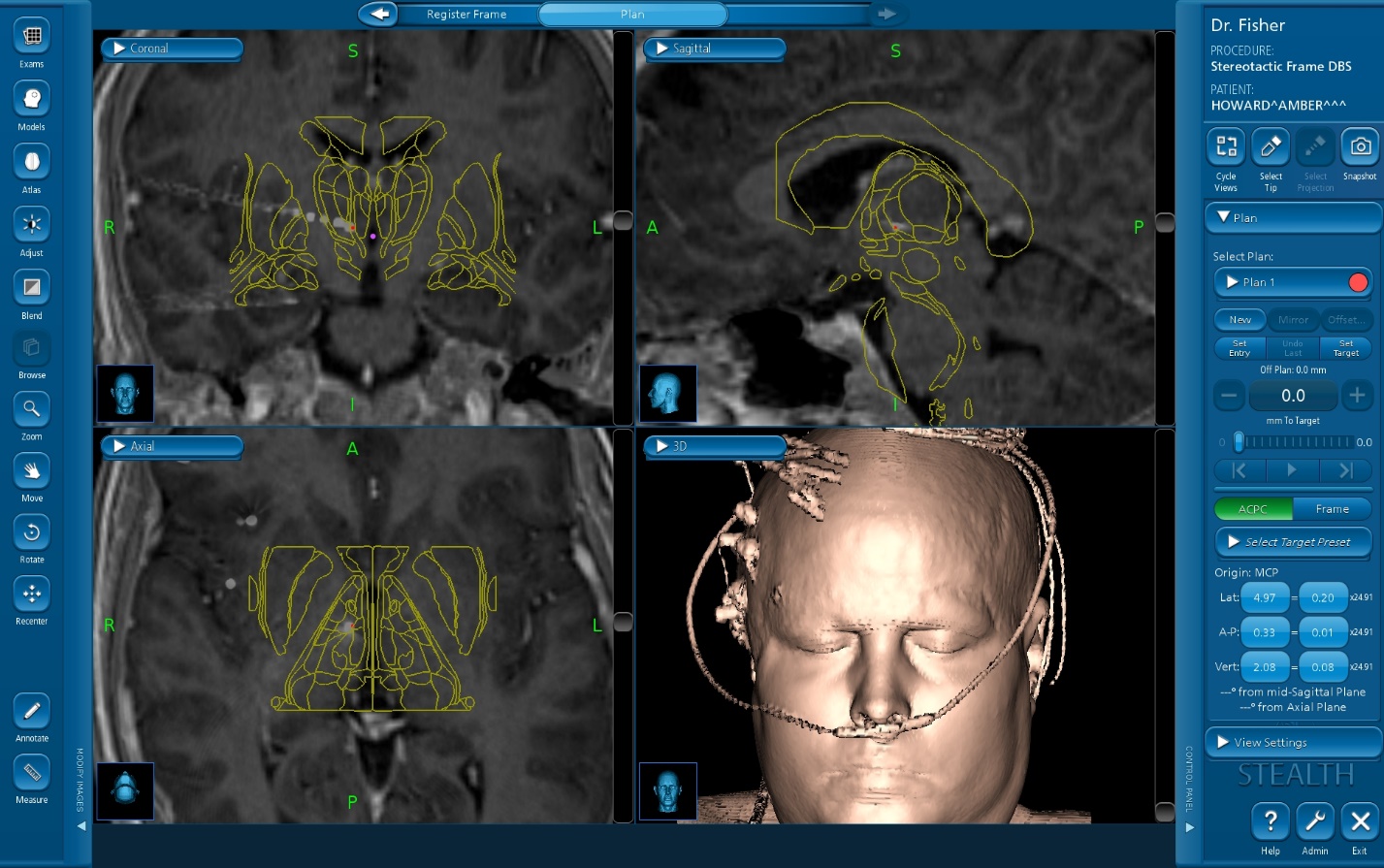

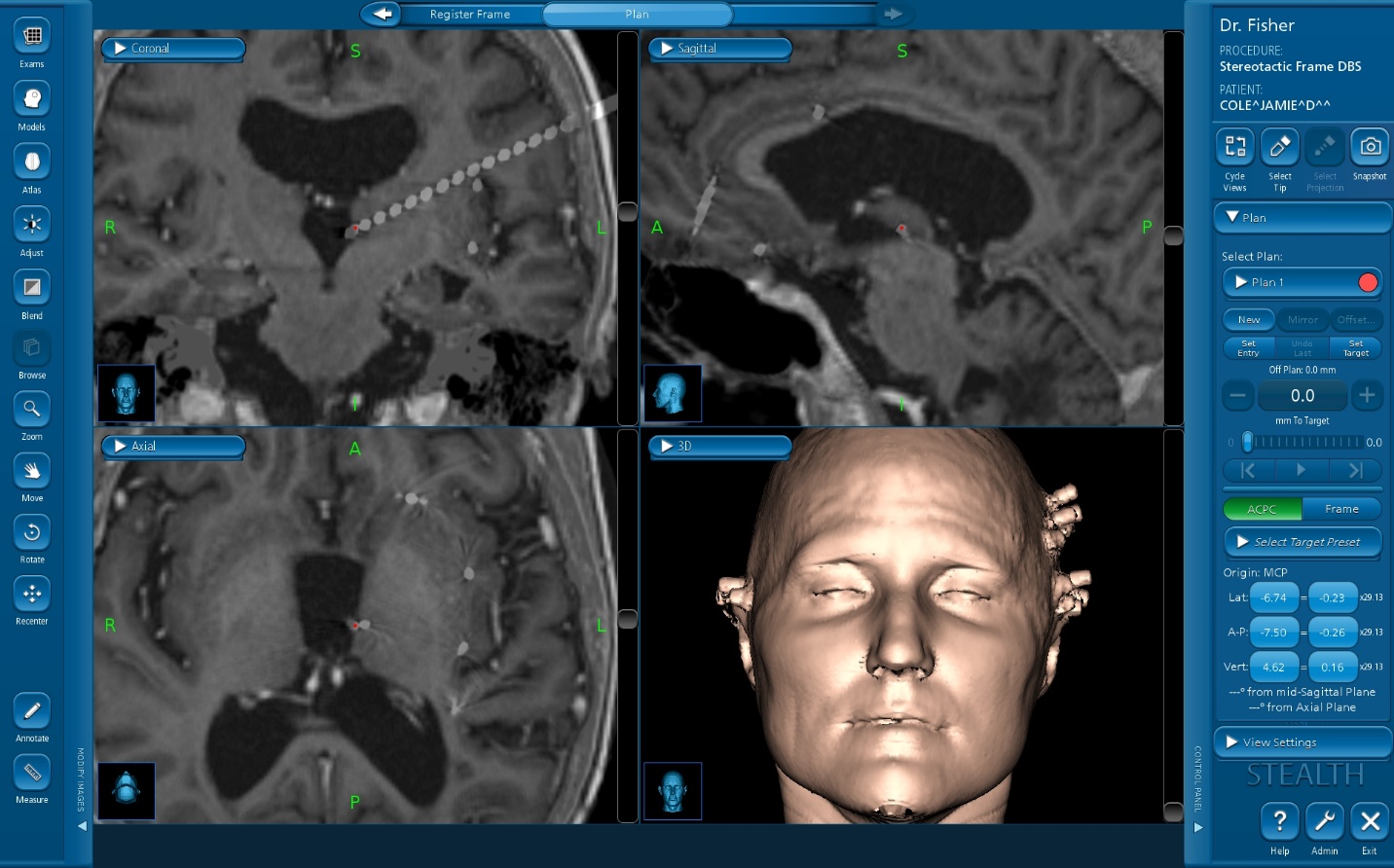
